# Unravelling the virome in birch: RNA-Seq reveals a complex of known and novel viruses

**DOI:** 10.1101/740092

**Authors:** Artemis Rumbou, Thierry Candresse, Armelle Marais, Laurence Svanella-Dumas, Maria Landgraf, Susanne von Bargen, Carmen Büttner

## Abstract

High-throughput sequencing (HTS), combined with bioinformatics for *de novo* discovery and assembly of plant virus or viroid genome reads, has promoted the discovery of abundant novel DNA and RNA viruses and viroids. However, the elucidation of a viral population in a single plant is rarely reported. In five birch trees of German and Finnish origin exhibiting symptoms of birch leaf-roll disease (BRLD), we identified in total five viruses, among which three are novel. The number of identified virus variants in each transcriptome ranged from one to five. The novel species are genetically - fully or partially - characterized, they belong to the genera *Carlavirus, Idaeovirus* and *Capillovirus* and they are tentatively named *birch carlavirus*, *birch idaeovirus*, and *birch capillovirus*, respectively. The only virus systematically detected by HTS in symptomatic trees affected by the BRLD was the recently discovered birch leafroll-associated virus. The role of the new carlavirus in BLRD etiology seems at best weak, as it was detected only in one of three symptomatic trees. Continuing studies have to clarify the impact of the carlavirus to the BLRD. The role of the *Capillovirus* and the *Idaeovirus* within the BLRD complex and whether they influence plant vitality need to be investigated. Our study reveals the viral population in single birch trees and provides a comprehensive overview for the diversities of the viral communities they harbor.

## Introduction

Symptoms in birch trees (*Betula* sp.) caused by various viruses and related to the birch leaf-roll disease (BLRD) are observed throughout Europe [1–2]. Diseased birches exhibit foliar disorders including vein banding, leaf roll, mottling, necrotic lesions and tip dieback. Based on earlier studies on virus-diseased trees it is assumed that BLRD might significantly reduce the tree’s photosynthetic capacity and contribute to tree decline [3]. Due to lack of knowledge, risk analyses and prevention measures, the disease has effectively spread throughout Europe and has until now been reported in European countries with diverse climatic conditions such as Finland, Sweden, Norway, Germany, Austria, UK and France, including stands of a Mediterranean island [2–5].

The initial hypothesis for the BLRD disease etiology implicated *Cherry leaf roll virus* (CLRV) as the disease main causal agent [1–2; 4–5]. However, after long-lasting trials applying conventional viral detection methods (virus purification, RT-PCR, double-stranded-RNA isolation, virus mechanical transmission), this virus could not be convincingly associated with the appearance of disease symptoms. To identify BLRD etiology, a birch metagenomic study was therefore initiated. A RNA-Seq analysis revealed for the first time in diseased trees from Germany and Finland birch leafroll-associated virus (BLRaV), the first reverse-transcribing DNA virus (*Badnavirus*, *Caulimoviridae*) discovered in birches [6]. Due to the clear correlation between BLRaV presence and BLRD-related symptoms and because symptoms were reproduced after grafting healthy seedlings with scions from BLRaV-infected trees, this virus is now considered to be strongly associated with BLRD. However, apart from the novel badnavirus, the applied high-throughput sequencing (HTS) strategy enabled the characterization of the entire genetic information of the ensemble of viruses in the tested samples, providing for first time insights into the birch metavirome.

The wide application of HTS technologies has significantly facilitated the discovery and characterization of viral agents in trees. Several HTS-based approaches have been conducted to overcome traditional approaches, which has resulted in the identification of known and so far unknown viruses providing insight into the virome of a species. Regarding fruit trees, in the last five years HTS use has led to the discovery of many new viruses [7–12]. Similarly to the present work, RNA viromes are detected in six peach trees identifying up to six viruses and viroids in each tree [13]. HTS data in nectarine provided insight to the etiology of stem pitting disease [14]. Diseased grapevine plants infected by *Grapevine Pinot gris virus* are found based on HTS data to acquire very complex viromes [15]. The total RNA-Seq approach concretely in grapevine resulted in complete virus and viroid genomes through *de-novo* assembly [16–17]. Building on this success, the movement to apply HTS for routine virus detection is gaining momentum. However, as far as challenges for the HTS for virus detection are concerned, validation should be taken into consideration and could focus on minimizing the risk of false negative results [18].

In the present study a first look into the birch virome is attempted. The first results on the virome of five birch trees of German and Finnish origin are described, defined as the exhaustive collection of nucleic acid sequences deriving from viral agents. A complex of known and novel viruses - including the recently discovered BLRaV - and of diverse variants of those agents was found to infect the tested samples. Apart from the already described viruses, novel species from the genera *Carlavirus, Idaeovirus* and *Capillovirus* are here identified and - fully or partially - genetically characterized.

## Materials and methods

### RNA-Seq and sequence assembly

Two twigs originating from a *Betula pubescens* donor tree (Bpub3) with severe BLRD leaf symptoms (vein banding, leaf chlorosis and necrosis, leaf rolling) from Rovaniemi (Finland) were grafted on two non-symptomatic *B. pubescens* rootstocks, generating grafted seedlings BpubFin407501_3A and BpubFin407507_3I. One twig originating from a *B. pendula* tree (Bpen 5) from Berlin, Germany also exhibiting BLRD symptoms was grafted on a non-symptomatic *B. pendula* rootstock, generating grafted seedling BpenGer407526_5M. Two CLRV-negative and symptomless birch seedlings of respectively *B. pubescens* (BpubGerNo4) and *B. pendula* (BpenGerM0197542), obtained from the same German nursery were used as negative controls. As rootstocks for the grafting and as control trees, two-year-old sprouting birches were used (nursery Reinke GbR Baumschulen, Rellingen, Germany). The following growing season, symptoms similar to the ones exhibited by the donor trees could be observed on the grafted birches already at the beginning of May and developed further until the end of September (Fig 1). No symptoms were observed on the negative, symptomless control trees.

**Fig 1.**
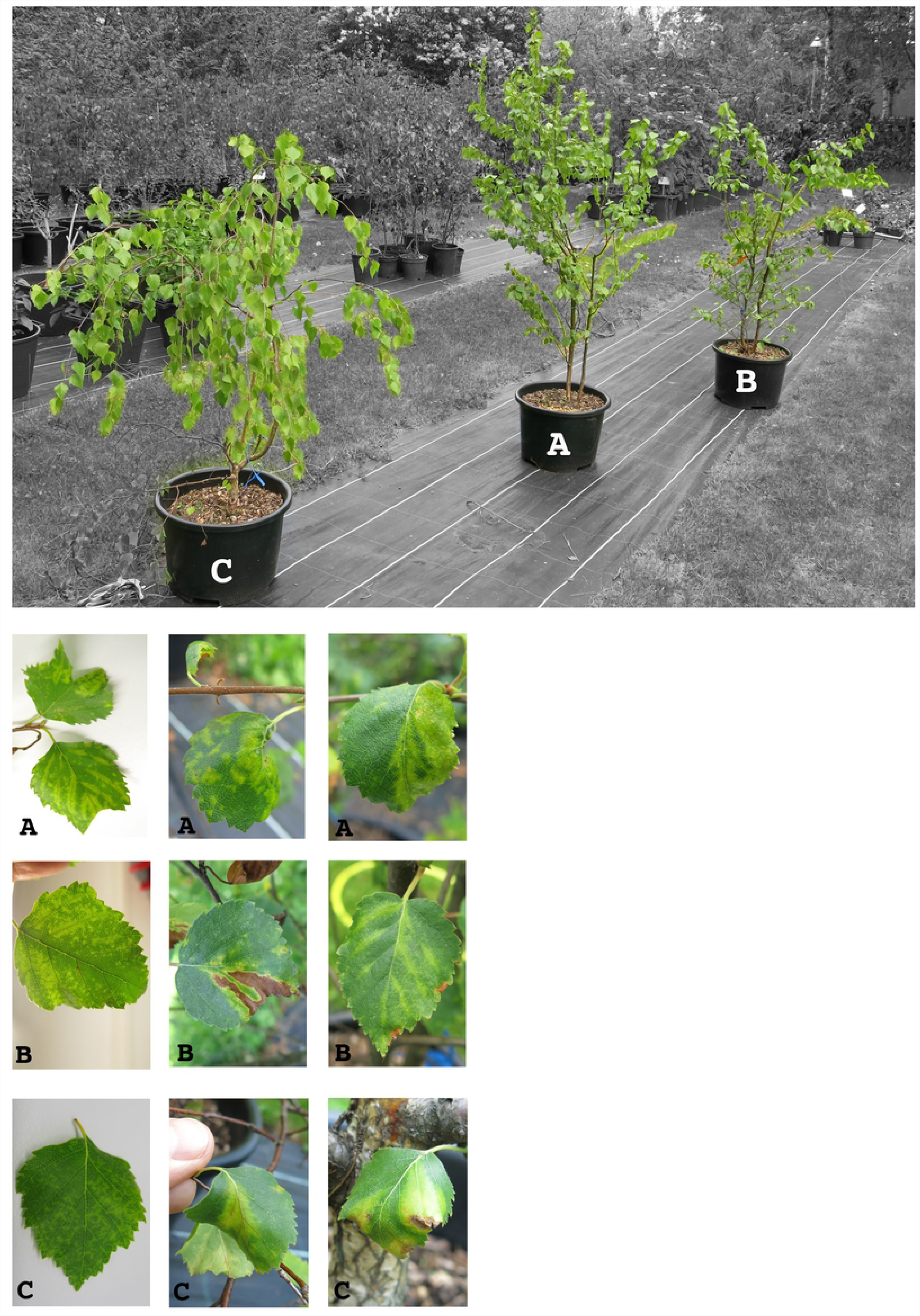
Leaf symptoms exhibited on the grafted birch seedlings BpubFin407501_3A (A), BpubFin407507_3I (B) and BpenGer407526_5M (C). Symptoms on seedling A (from left to right): Chlorotic vein banding, mottling, vein banding and leaf-roll. Symptoms on seedling B: mottling, leaf necrosis (final stage developed from chlorosis), vein chlorosis. Symptoms on seedling C: mottling, leaf roll and vein banding, vein banding with necrosis and leaf roll.

In 2014, pooled samples of five symptomatic leaves randomly selected from the seedlings canopy were used for RNA extraction. Similar leaf pools obtained from the symptomless trees were used in parallel. Total RNAs isolation, cDNA synthesis and preparation for RNA-Seq analysis with the Illumina HiSeq2500 system are fully described in Rumbou et al., 2018 [6]. 100 bp-long paired-end sequence reads corresponding to 50-100 Mb data/sample were generated. All HTS data processing and analysis were performed using CLC Genomics Workbench version 7.0.4. Reads were first submitted to quality filtering and trimming. The resulting cleaned reads were then assembled into contigs that were finally annotated by BlastN and BlastX against the GenBank database.

### Taxonomic analysis of the metagenome

The taxonomic content of the obtained datasets, as provided by the Blast analyses was visualized using MEGAN [19], in which the result of the Blast analyses are parsed to assign the best hits to appropriate taxa in the NCBI taxonomy. As a result, the taxonomical content (“species profile”) of the sample from which the reads were collected was estimated, with a particular focus on viral species (Fig 2; A-E).

**Fig 2.**
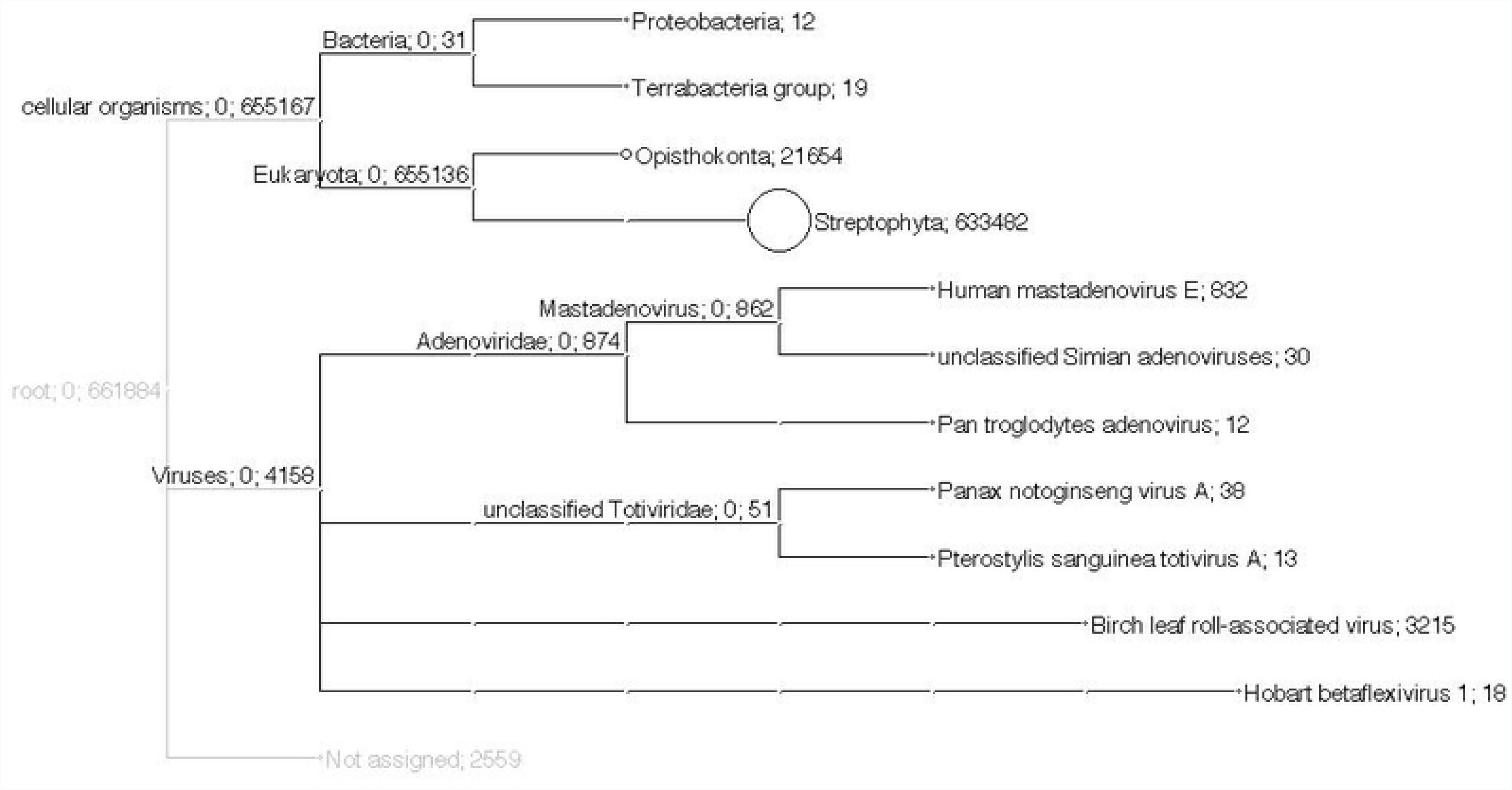
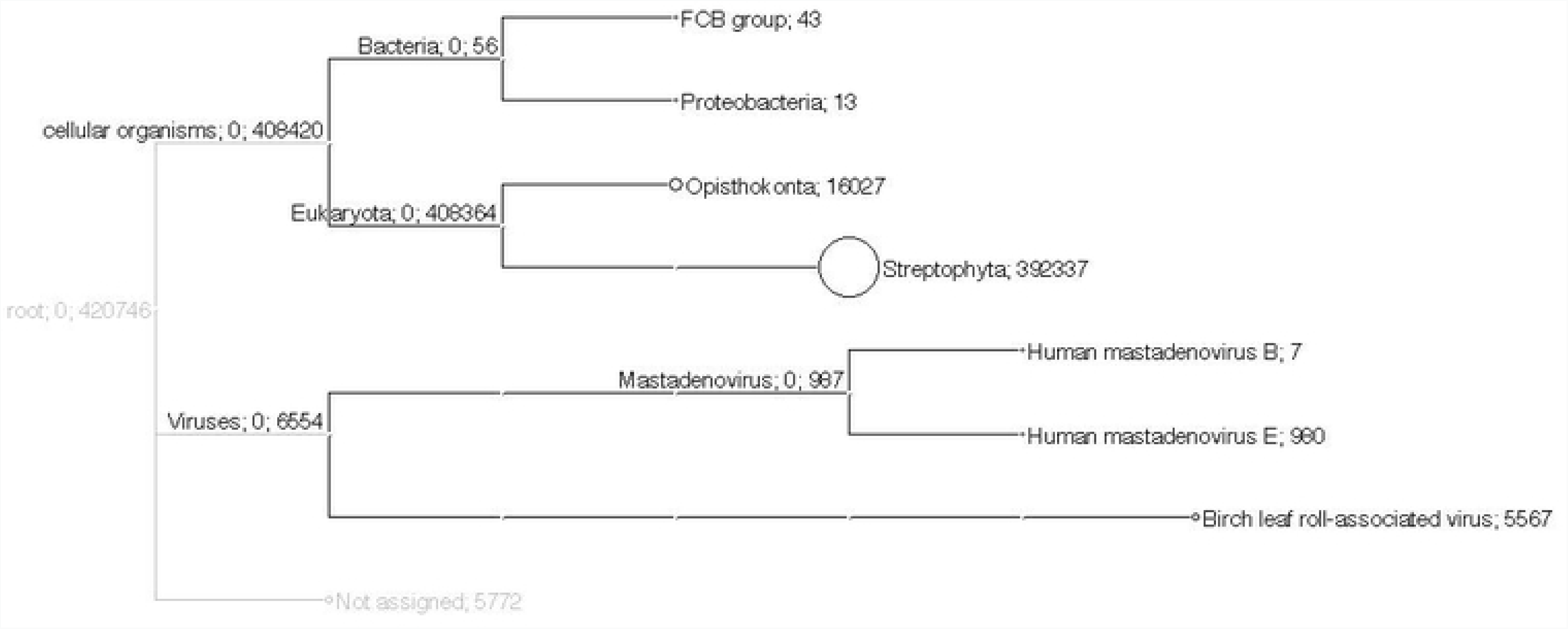
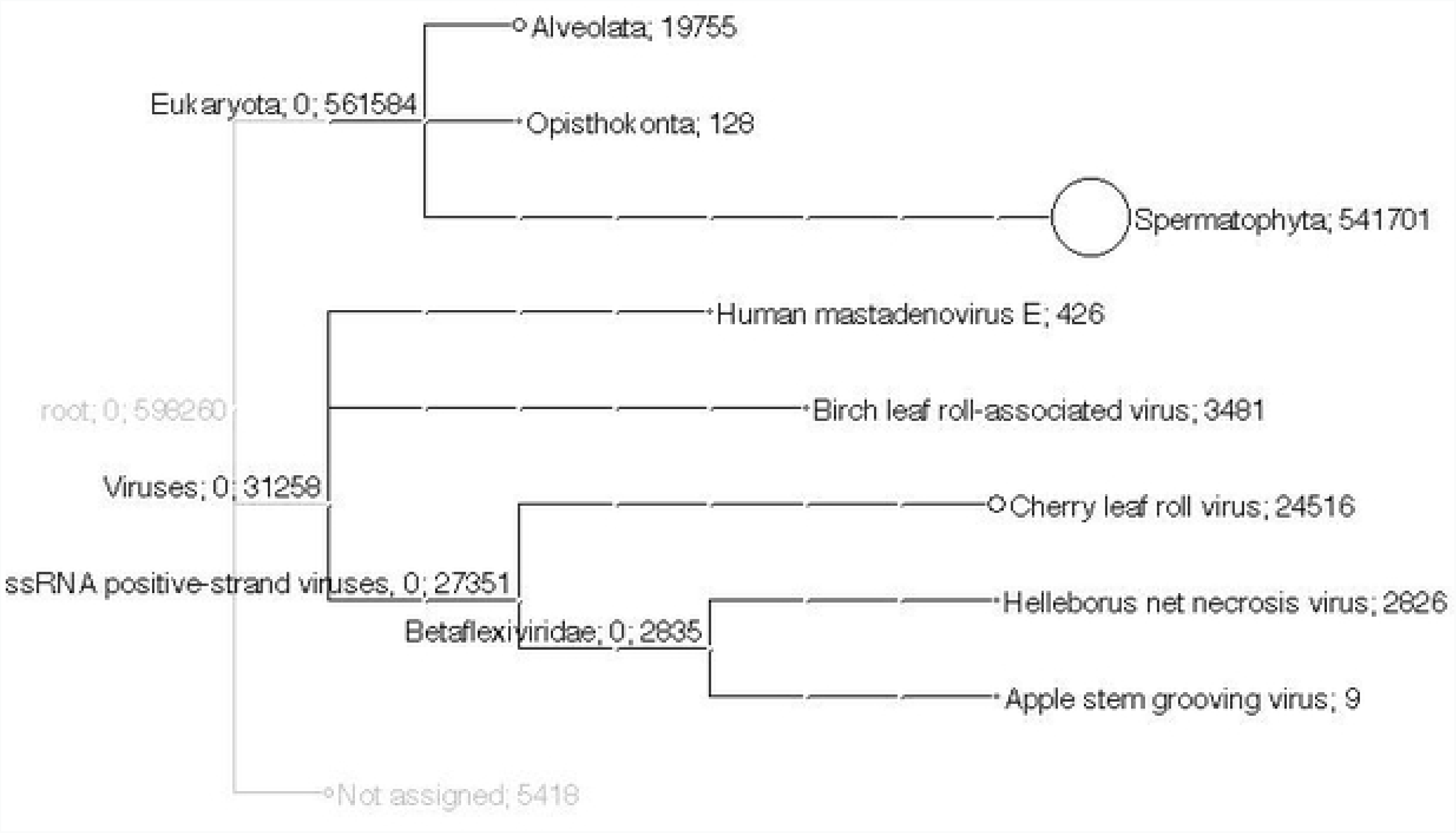
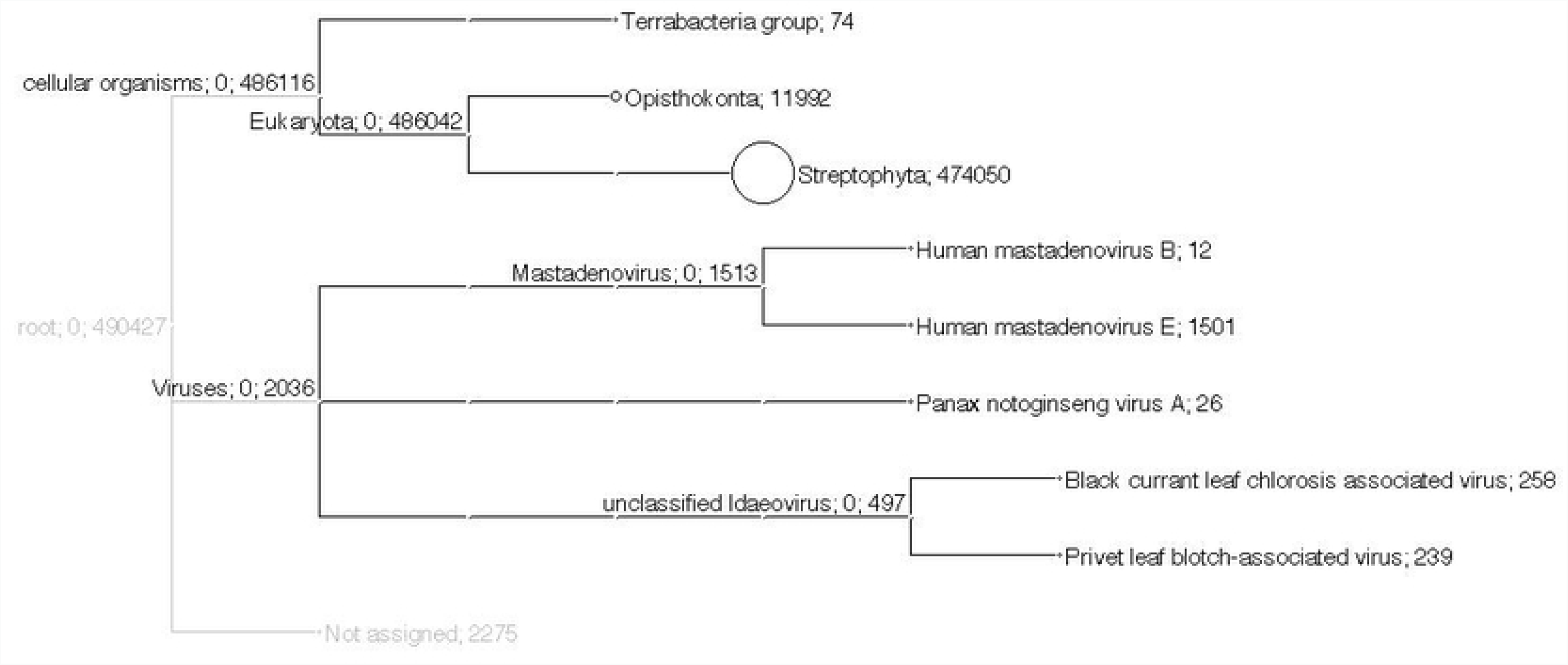
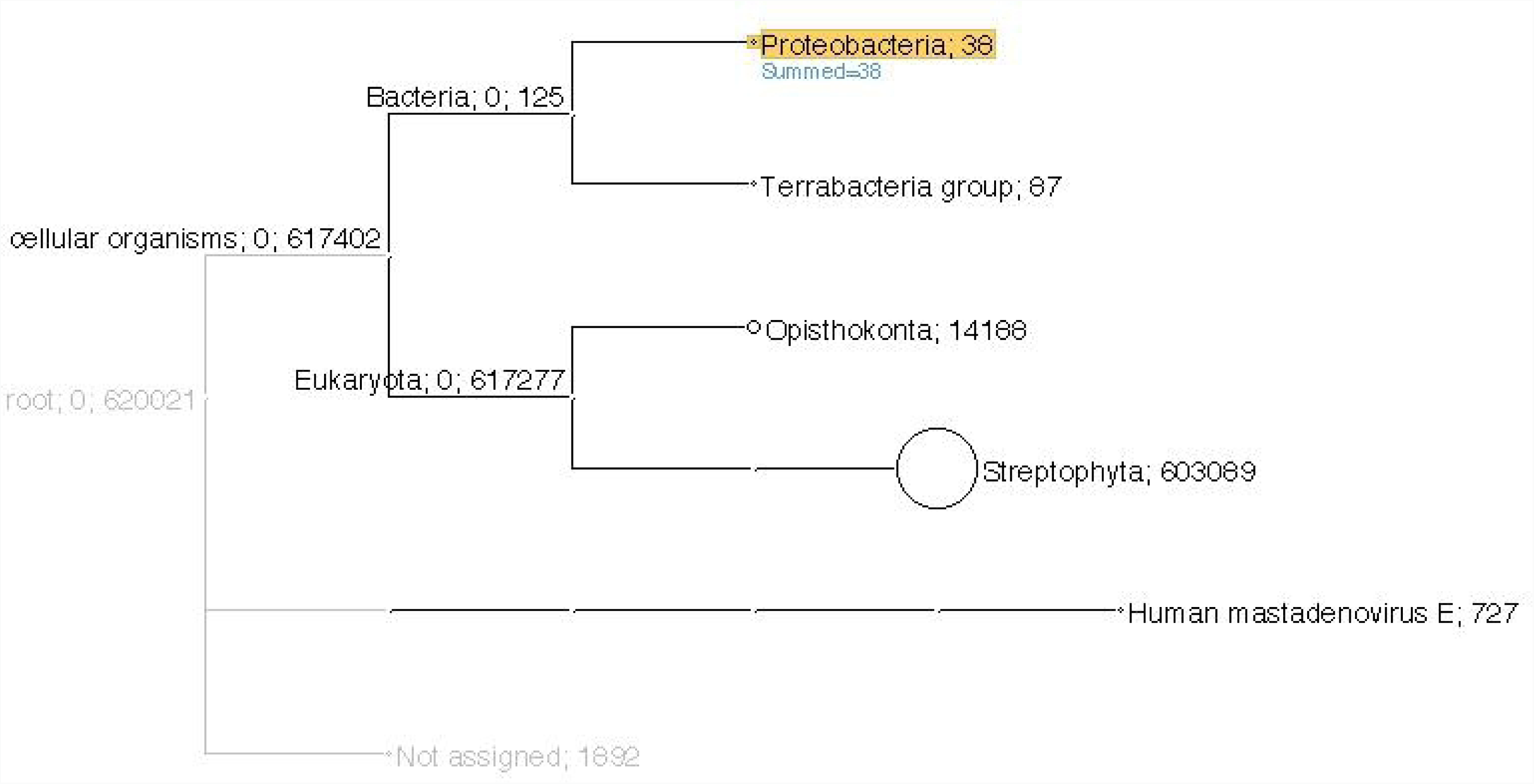
Taxonomical content of the birch samples analyzed by RNA-Seq with focus on the virome. A. symptomatic birch BpubFinn407501_3A, B. symptomatic birch BpubFinn407507_3I, C. symptomatic birch BpenGer407526_B5, D. symptomless birch BpubGer4 and E. symptomless birch BpenGerMO197542. Labels include taxon; number of reads assigned to taxon, summarized number of reads.

### Validation of the presence of novel viruses in birches

In order to confirm the presence of the identified novel viruses, specific RT-PCR assays were performed using virus-specific primers designed using the sequence of the scaffolds assembled for each agent (Table 1). These primer pairs were designed using OligoCalc [20] and respectively target regions within the RdRp domain (Carla for/rev; nt 926 – 1558) for the new carlavirus, within the methyl transferase (MTR) domain (RNA1) for the new idaeovirus (Idaeo_for/rev; nt 487 – 1060) and within the coat protein domain for the new capillovirus (Betaflexi_for/rev; nt 95 - 552).

**Table 1.**
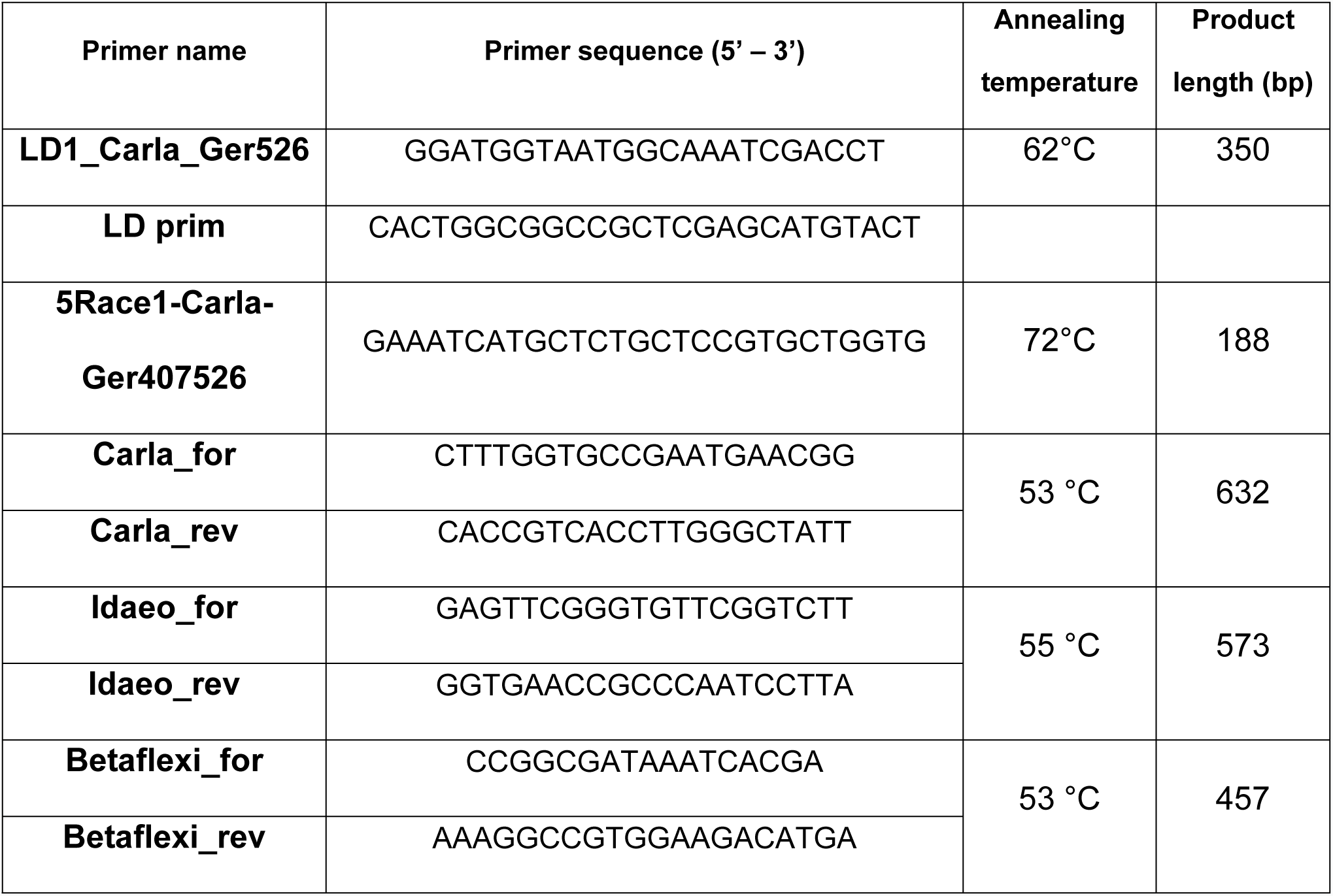
Primers used for genome completion and for the specific detection of the novel viruses.

Pooled samples of 3 to 5 leaves from different twigs of each tree were used. The first strand cDNAs were synthesized from 1 µg of total RNA in a 20 µl reaction volume of 1 x RT buffer (Thermo Scientific) containing 1 µM dNTPs mix, 200 U RevertAid Premium reverse transcriptase (Thermo Scientific), 20 U Ribolock RNase inhibitor (Thermo Scientific) and 100 pmol of random hexamer-oligonucleotides (Biomers.net GmbH). Subsequent PCR amplifications were conducted in a 25 µl volume of 1 x DreamTaq Buffer (Thermo Scientific) containing 0.2 µM dNTP mix, 0.625 U of DreamTaq DNA polymerase and 1 µM of each forward and reverse primer (Table 1). The thermal cycles were as follows: 2 min at 95 °C followed by 35 cycles at 95 °C for 30 s, T_anneal_ for 30 s, 72 °C for 40 s, with a final extension step of 72 °C for 5 min. Omitting the primers sequences, the amplified fragments are 592 nucleotides (nt) long for the carlavirus, 533 nt for the idaeovirus and 420 nt for the capillovirus. PCR products were directly submitted for Sanger sequencing (Macrogen) without previous cloning.

### Completion of the carlavirus genome ends

Assuming a dsRNA stage of the tentative carlavirus, 5’ and 3’ ends of the genome were determined using a 5’ Rapid amplification of cDNA-ends (5’ RACE) strategy, and a polyA-anchored Long Distance-RT-PCR, respectively. The 5’RACE reaction was performed according to the kit manufacturer’s instructions (Clontech / Ozyme, Saint-Quentin en Yvelines, France) (Tprimer 5Race1-Carla-Ger407526; Table 1), and the 3’ genome end was amplified following the protocol described by Youssef et al. [21] (primers LD1_Carla_Ger526 and LD prim; Table 1).

### Phylogenetic analyses of Carlavirus sequences

Multiple nucleotide or amino acid sequence alignments were performed as well as pairwise sequence identity calculations using AliView version 1.17.1 [22]. For the phylogenetic comparisons of complete RdRp and MP regions, all identified carlavirus species represented in GenBank to date were used. Bootstrapped Maximum likelihood (ML) trees were constructed with MEGA6 [23]. Robustness of nodes of the phylogenetic tree was assessed from 1,000 bootstrap resamplings, and values > 70% displayed for trees internal nodes.

## Results

### Birch metagenome taxonomic analysis with focus on the virome

The results obtained by MEGAN analysis regarding the taxonomic content of contigs assembled from the RNA-Seq reads are shown in Fig 2 (A - E), together with the number of reads assigned to each taxon. For symptomatic sample BpenGer407526-B5, out of the 598.260 reads assessed, 561.584 belong to Eucaryota, most of them to the Phylum *Spermatophyta*, where *Betula* sp. is classified and the rest to Protista (Alveolata) and Opisthokonta (Fungi and Vertebrata) (Fig 2, C). From the 31.258 viral reads, 3.481 reads are attributed to birch leaf roll-associated virus (*Badnavirus*, *Caulimoviridae*) and specifically to two variants of this virus (see ref. 6 for detailed description). Within the single-stranded RNA viruses, 24.516 reads belong to cherry leaf roll virus (*Nepovirus*, *Secoviridae*) while 2.835 reads are analyzed as representing agent(s) in the family *Betaflexiviridae*, with affinities to helleborus net necrosis virus (2.826 reads) and apple stem grooving virus (9 reads). The presence of 426 reads from Human mastadenovirus E (dsDNA viruses) is attributed to possible contamination of the sample during sample handlings or sequencing (all 5 samples exhibit presence of this human virus)

*B. pubescens* samples BpubFinn407501-3A and BpubFinn407507-3I are both found to be infected by the badnavirus BLRaV with a high number of reads (Fig 2; A and B). Furthermore, in the sample BpubFinn407501-3A, 18 reads are attributed to hobart betaflexivirus 1, an unclassified member of the *Betaflexiviridae* family, and 51 reads to Totiviruses, known to infect fungi (Fig 2; A). The non-symptomatic birch seedlings are negative for all viruses present in the symptomatic ones. However, 497 reads in the sample BpendGerMO197542 are attributed to the genus *Idaeovirus*, with closest relatives identified as black currant leaf chlorosis-associated virus and privet leaf blotch-associated virus (Fig 2; E).

An overview of the obtained RNAseq data identified in each sample is presented in Table 2.

**Table 2.** Virome data generated for each birch seedling. The number of reads and their percentage in the sample as well as the validation output through RT-PCR (+/-) are presented. (BLRaV: birch leafroll-associated virus; CLRV: cherry leaf roll virus; BiCV: birch carlavirus; BCV: birch capillovirus; BIV: birch idaeovirus).

|  | Bpub Finn<br>407501_3A |  |  | Bpub Finn<br>407507_3I |  |  | BpenGer<br>407526_B5 |  |  | Bpen Ger<br>MO197542 |  |  | Bpub<br>Ger4 |  |  |
| --- | --- | --- | --- | --- | --- | --- | --- | --- | --- | --- | --- | --- | --- | --- | --- |
|  | Number of<br>reads | % | PCR | Number of<br>reads | % | PCR | Number of<br>reads | % | PCR | Number of<br>reads | % | PCR | Number of<br>reads | % | PCR |
| **total** | 803.120 |  |  | 613.923 |  |  | 725.231 |  |  | 546.722 |  |  | 682.408 |  |  |
| **BLRaV** | 3211 | 0.4 | + | 5567 | 0.9 | + | 3529 | 0.49 | + | 1 | 0.0002 | - | 1 | 0.00015 | - |
| **CLRV** | 3 | 0.0004 | + | 3 | 0.0005 | + | 10896 | 1.5 | + | 0 | 0 | - | 2 | 0.0003 | - |
| **BiCV** | 3 | 0.0004 | - | 5 | 0.0008 | - | 2881 | 0.397 | + | 2 | 0.00037 | - | 1 | 0.00015 | - |
| **BCV** | 21 | 0.0026 | + | 3 | 0.0005 | + | 20 | 0.003 | + | 5 | 0.0009 | + | 2 | 0.00029 | + |
| **BIV** | 0 | 0 | - | 0 | 0 | - | 0 | 0 | - | 195 | 0.036 | + | 0 | 0 | - |

### Full genome assembly of a new birch CLRV variant

CLRV was only detected in one of the tested symptomatic birches, the *B. pendula* BpenGer407526_B5 from Berlin. The full-length genome, which consists of two RNA segments, was assembled. RNA1 is 7,848bp-long and highly similar to the birch isolate already deposited in GenBank (LT883167, 96 % nt identity). RNA2 is 6,459bp-long and exhibits a lower level of identity with the birch CLRV isolate (LT883166, 91 % nt identity), similar to what is observed with the cherry CLRV isolate (JN104385, 91 % nt identity). The genomic sequences of this new CLRV variant have been deposited in GenBank under accession numbers MK402281 (RNA1) and MK402282 (RNA2).

### Partial genome assembly of a novel idaeovirus

In the dataset from the symptomless seedling BpenGerMO197542, two long contigs of an uncharacterized virus with affinities to idaeoviruses were assembled (see Fig 2; E). The first contig is 5,232bp-long and encodes a putative protein, which in the BLAST analysis shows high level of homology with the ORF1 of the black currant leaf chlorosis associated virus (BCLCaV, YP_009361854, 63 % aa identity) a novel, recently described idaeovirus (James and Phelan, 2017). The ORF1 initiates at nt position 283-285 of the contig and codes for a 1649 aa putative replication-associated protein with conserved methyltransferase (MTR), helicase (HEL) and RNA-dependent RNA polymerase (RdRp) domains. However, the protein is not complete as a stop codon is not reached and amino acids of the Cter end of the protein are missing compared to BCLCaV. The second contig is 1,595bp-long and harbours 2 ORFs. The first ORF initiates at nt position 5-7 of the contig and ends at positions 1091-1093, and encodes a putative 362 aa-long movement protein exhibiting homologies with the corresponding protein of BCLCaV (YP_009361835, 43% aa identity). The second ORF of the RNA2 initiates at nt positions 1090-1092, overlapping with the stop codon of the first ORF as is also observed for BCLCaV [24]. This second ORF encodes a putative coat protein (CP), which is homologous with the CP of BCLCaV (YP_009361836, 47% aa identity), however only the first 167 aa of the protein are available, as the genome is not completely covered by the contig. The presence of this novel virus was validated by RT-PCR with specific primers (Idaeo_for/Idaeo_rev; Table 1) in the tested seedling. Sequencing of the amplified products provided a sequence identical with the original contig thus further confirming the infection. We suggest, therefore, that the two contigs correspond to a novel idaeovirus, tentatively named as birch idaeovirus (BIV). The obtained incomplete viral sequences were deposited in GenBank under accession numbers MK402235 (RNA1) and MK402236 (RNA2).

In August 2015 the symptomless seedling in which BIV was detected developed sporadic virus-like symptoms of variegation in a few leaves (Fig 3). Given the presence of the new ideaovirus in the plant, the possibility that this virus is responsible for these symptoms should be further investigated.

**Fig 3.**
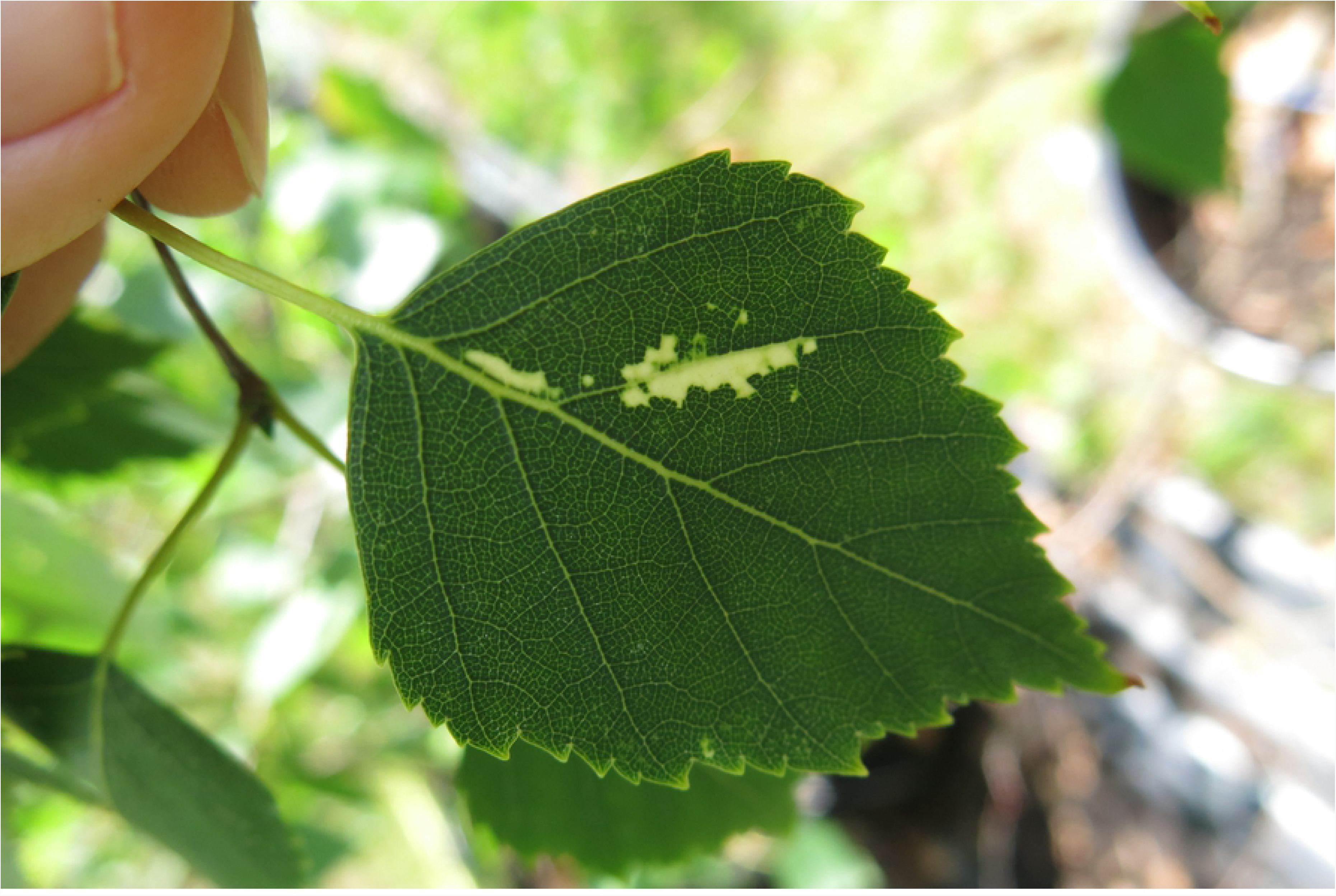
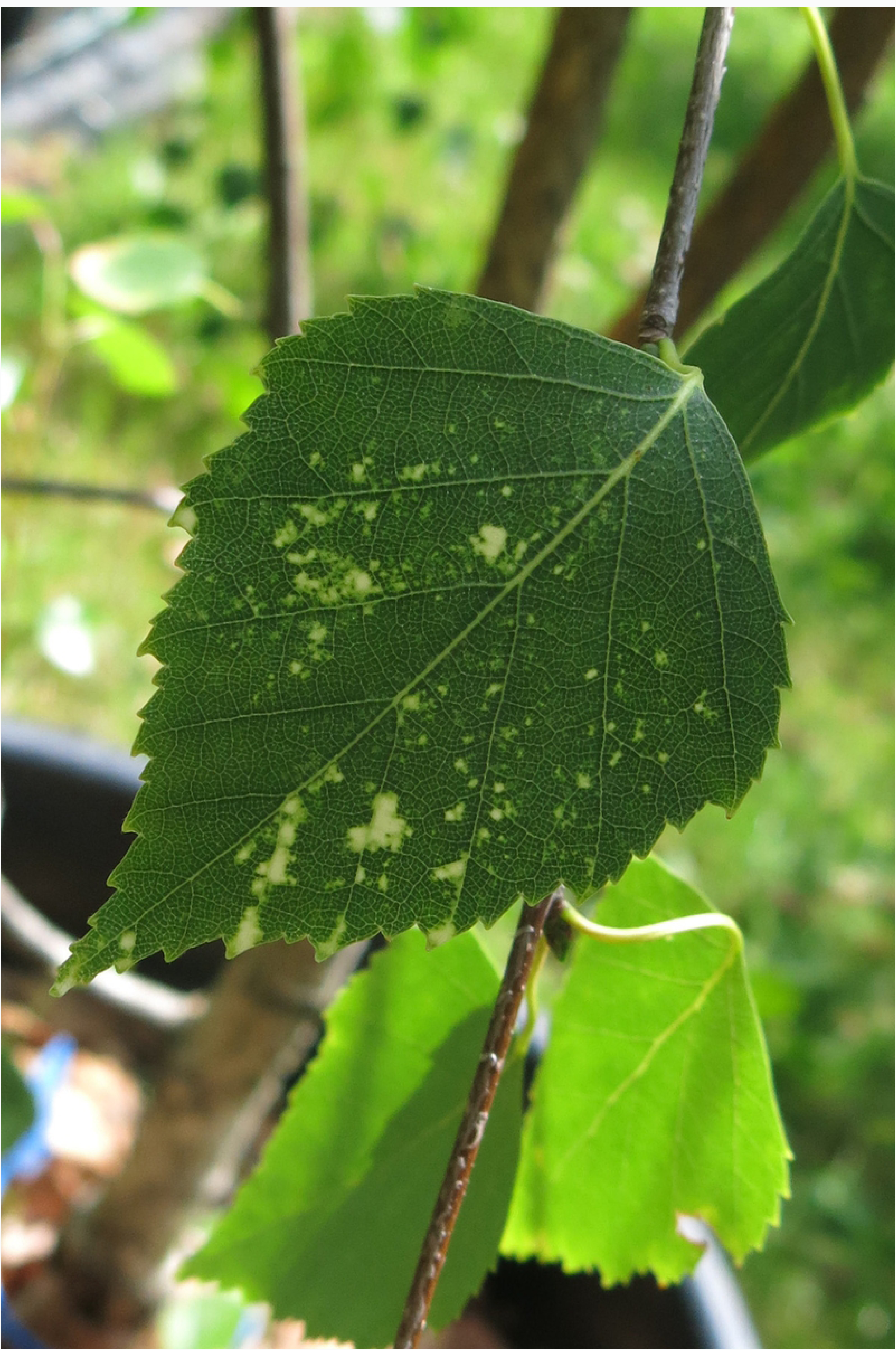
Symptoms appeared in the seedling BpenGerMO197542.

### Assembly of a capillo-like virus sequence

From the RNAseq dataset of the BpenGer407526_B5 seedling a 821bp-long contig was assembled, which exhibits homologies with apple stem grooving virus (ASGV, *Betaflexiviridae, Capillovirus*, Fig 2; C). Further attempts to obtain longer sequences of the novel virus by means of PCR resulted in a 1114bp-long contig extending all the way to the 3’-poly(A) tail. This contig was submitted to GenBank under accession number MK402233. Within the contig, the 250 aa-long (753 nt) coat protein sequence of this novel virus (nt positions 60-812) is encoded. In the BLASTP analysis this putative protein shares low but significant identity with the CP of ASGV (AFH75121, 30% identity). Furthermore, a 597bp-long sequence covering part of the same genomic region was assembled from the BpubFinn407501_3A reads (accession number MK402234), showing 98,7% nt identity with the first one. This contig is assembled from the reads attributed to hobart betaflexivirus 1 in the Megan analysis of Fig 2; A.

As the encoded proteins of the new virus show less than 80% aa identity with CP sequences from other capilloviruses, they are suggested to represent a novel species of the genus *Capillovirus*. To investigate the assumption that the new virus is closely related to other capiloviruses, the phylogenetic relationships of the CP protein sequences from members of the *Betaflexiviridae* family were analyzed. In the obtained ML and NJ trees, the new virus reliably clustered within the capilloviruses clade (Fig 4). Concluding, the low amino acid identity with members of *Capillovirus* as well as the phylogeny generated for the CP regions suggest that this virus belongs to the genus Capilovirus and is therefore is tentatively named birch capillovirus (BCV).

**Fig 4.**
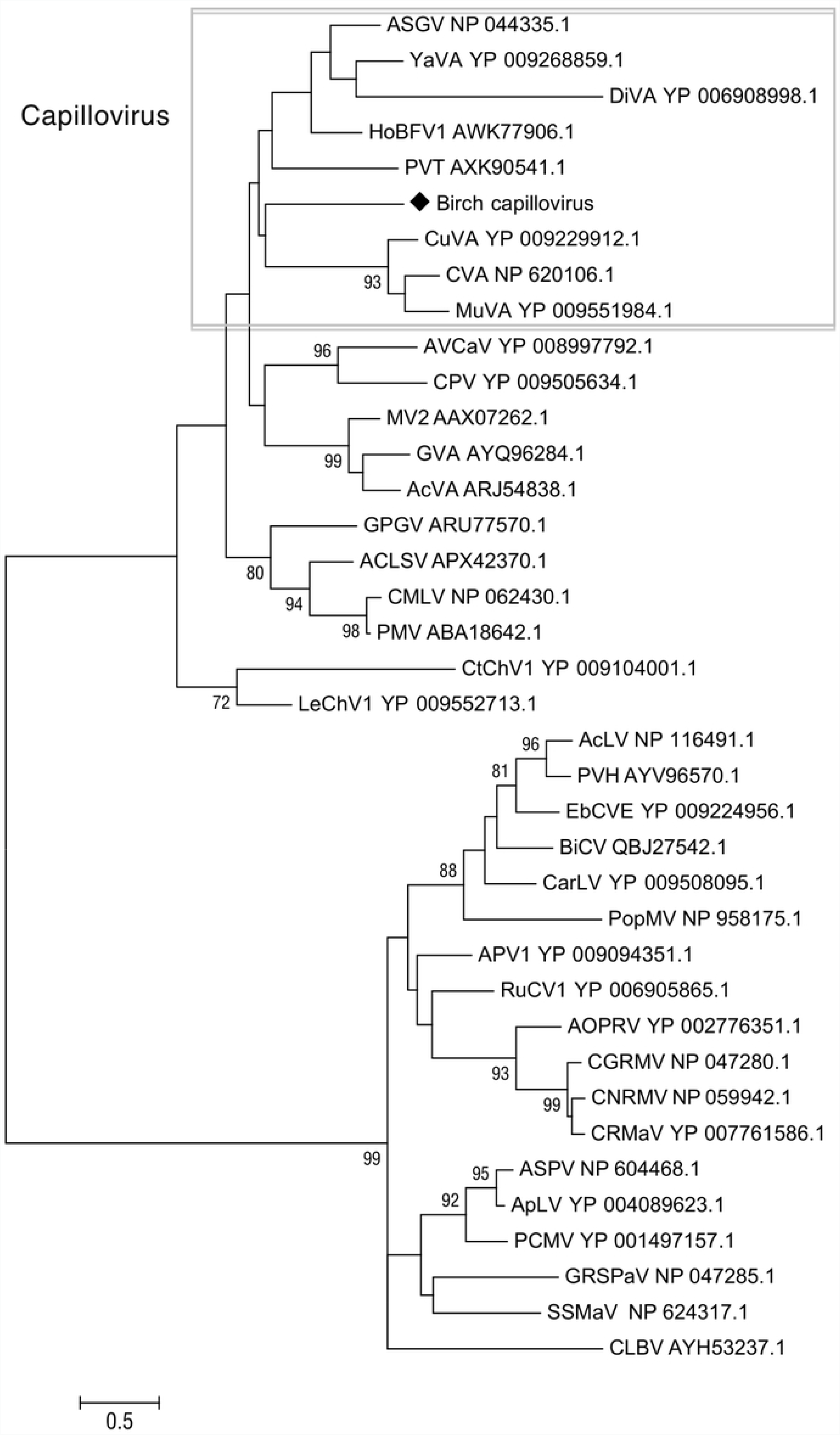
Phylogenetic tree reconstructed using the amino acid sequences of the CP of *Betaflexiviridae* members. The tree was reconstructed using the Maximum Likelihood method and the statistical significance of branches was evaluated by bootstrap analysis (1,000 replicates). Only bootstrap values above 70% are indicated. The scale bar represents 5% amino acid divergence. Members of the Capillovirus are indicated within the rectangle. Virus abbreviations and accession numbers are as follows: apple stem grooving virus (ASGV), yacon virus A (YaVA),diuris virus A (DiVA), hobart betaflexivirus 1 (HoBFV1), currant virus A (CuVA), cherry virus A (CVA), mume virus A (MuVA), potato virus T (PVT), apricot vein clearing associated virus (AVCaV), caucasus prunus virus (CPV), mint virus 2 (MV2), grapevine virus A (GVA), actinidia virus A (AcVA), grapevine Pinot gris virus (GPGV), apple chlorotic leaf spot virus (ACLSV), cherry mottle leaf virus (CMLV), peach mosaic virus (PMV), carrot Ch virus 1 (CtChV1), lettuce Chordovirus 1 (LeChV1), aconitum latent virus (AcLV), potato virus H (PVH), elderberry carlavirus E (EbCVE), birch carlavirus (BiCV), carnation latent virus (CarLV), poplar mosaic virus (PopMV), asian prunus virus 1 (APV1), rubus canadensis virus 1 (RuCV1), african oil palm ringspot virus (AOPRV), cherry green ring mottle virus (CGRMV), cherry necrotic rusty mottle virus (CNRMV), cherry rusty mottle-associated virus (CRMaV), apple stem pitting virus (ASPV), apricot latent virus (ApLV), peach chlorotic mottle virus (PCMV), grapevine stem pitting-associated virus (GRSPaV), sugarcane striate mosaic-associated virus (SSMaV), citrus leaf blotch virus (CLBV).

BCV presence was confirmed by RT-PCR not only in the seedling Bpen5MGer407526_B5 from which it originated, but also in all four other seedlings analyzed here (symptomatic and not symptomatic) and in other trees from Berlin and Rovaniemi (data not shown).

### Full genome assembly of a novel carlavirus from birch

BLASTN and BLASTX annotation of the assembled contigs from symptomatic birch BpenGer407526_B5 revealed one large contig exhibiting high BLAST scores with members of the genus *Carlavirus* (*Betaflexiviridae*) (see Fig 2; C, reads attributed by MEGAN to helleborus net necrosis virus). This 8,846 nt contig covers a near complete carlaviral genome, missing only the ends.

PolyA-anchored long-Distance (LD)-PCR and 5’RACE allowed the completion of the genome by generating sequences that perfectly matched the contig in the overlap regions. The presence of the virus was confirmed by specific RT-PCR performed in the seedling were it was firstly detected (BpenGer407526_B5) and in other trees in Berlin (data not shown). The full-length length genomic sequence of this novel agent was deposited in GenBank under accession number MH536506.

The genome of this virus is 8,896 base pairs (bp) long, which is close to the genome size of typical carlaviruses (8,3 – 8,7 kb) [25]. It shows a typical Carlavirus organization with 6 ORFs including a RNA-dependent RNA polymerase (RdRp; nt 61 – 6,084), three triple gene block proteins (TGB1; nt 6,153 – 6,857, TGB2; nt 6,835 – 7,167, TGB3; nt 7,169 – 7,381), a coat protein (CP; nt 7,432 – 8,442) and a nucleic-acids binding protein (NABP, ORF6; nt 8,442 – 8,843) (Fig 5). In the BLASTP analysis, the RdRp shows homologies with the corresponding protein of Carlaviruses, the closest being helleborus net necrosis virus (47% identity). The CP protein also shows significant levels of aa identity (41-55%) with other carlaviral CPs, the closest being elderberry carlavirus B (55% identity). The TGBs also have closest affinities to various carlaviruses: 53% identity for the TGB1 (elderberry carlavirus A), 54% for the TGB2 (poplar mosaic virus) and 71% for the TGB3 (carrot carlavirus) (Fig 4). The NABP encoded by ORF6 is closest to the corresponding protein of helleborus mosaic virus (42% identity).

**Fig 5.**
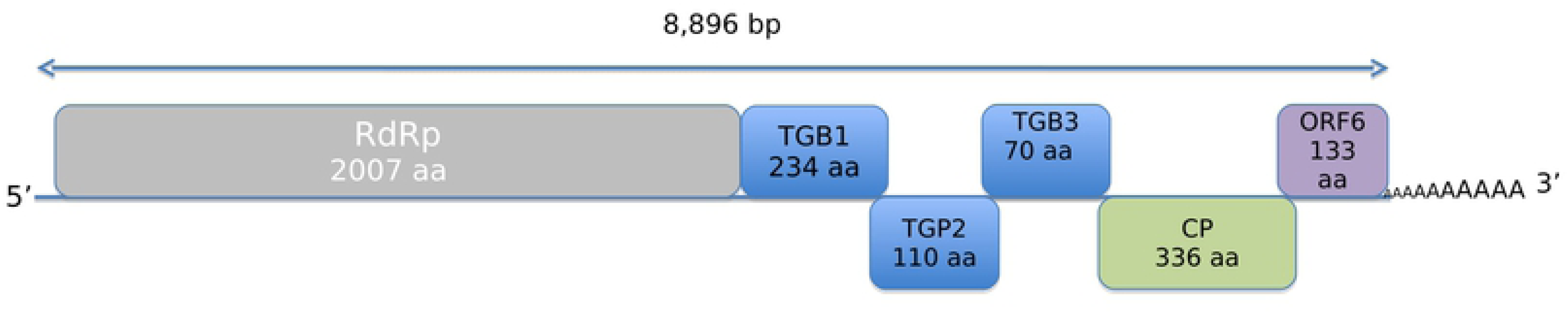
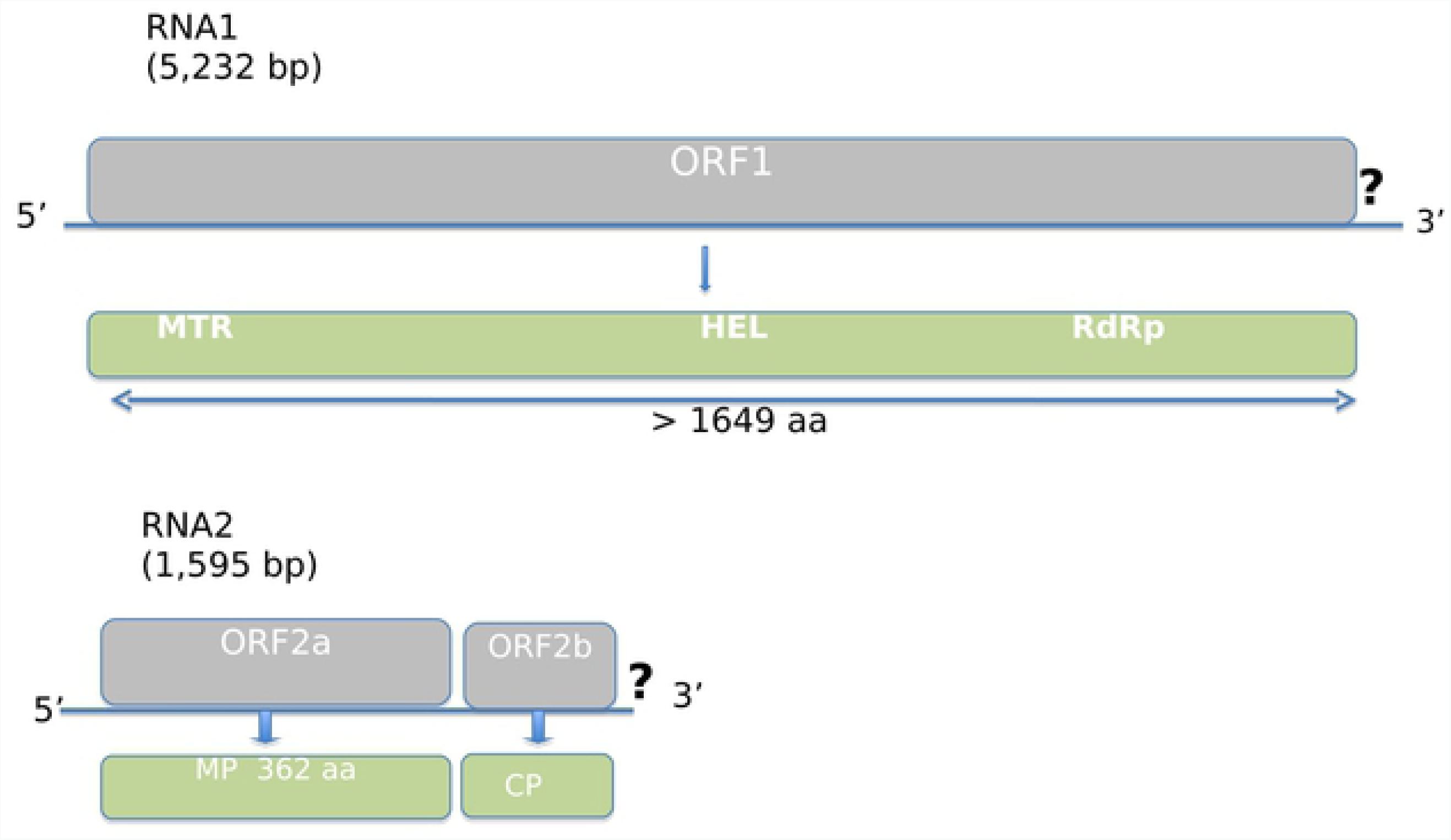
Schematic representation of the genome organization of the novel *Birch carlavirus* (BiCV; Fig 5A) and *Birch idaeovirus* (BIV; Fig 5B).

### Phylogenetic analysis of the novel carlavirus

Phylogenetic relationships between the birch carlavirus and the sequences of carlaviruses known to date were estimated, based on amino acid sequences comparisons. The topology of the trees was similar, irrespective of whether the ML or NJ algorithms were used. Fig 6 shows a representative ML tree obtained using the RdRp and CP protein sequences. The RdRp protein from the novel carlavirus from birch clusters together with the woody host carlaviruses poplar mosaic virus (PopMV) and elderberry carlavirus A, B and D (EBCVA, EBCVB, EBCD). The CP also clusters with the elderberry carlaviruses A, B and D (EBCVA, EBCVB, EBCD) and, more distantly, PopMV. The new virus is clearly only distantly related phylogenetically to all carlaviruses currently represented in the GenBank database (Fig 6), and exhibits less than 80% aa identity with the CP or RdRps of known carlaviruses. Taken together these results demonstrate that it represents a new member of the genus *Carlavirus* and it is, therefore, tentatively named birch carlavirus (BiCV).

**Fig 6.**
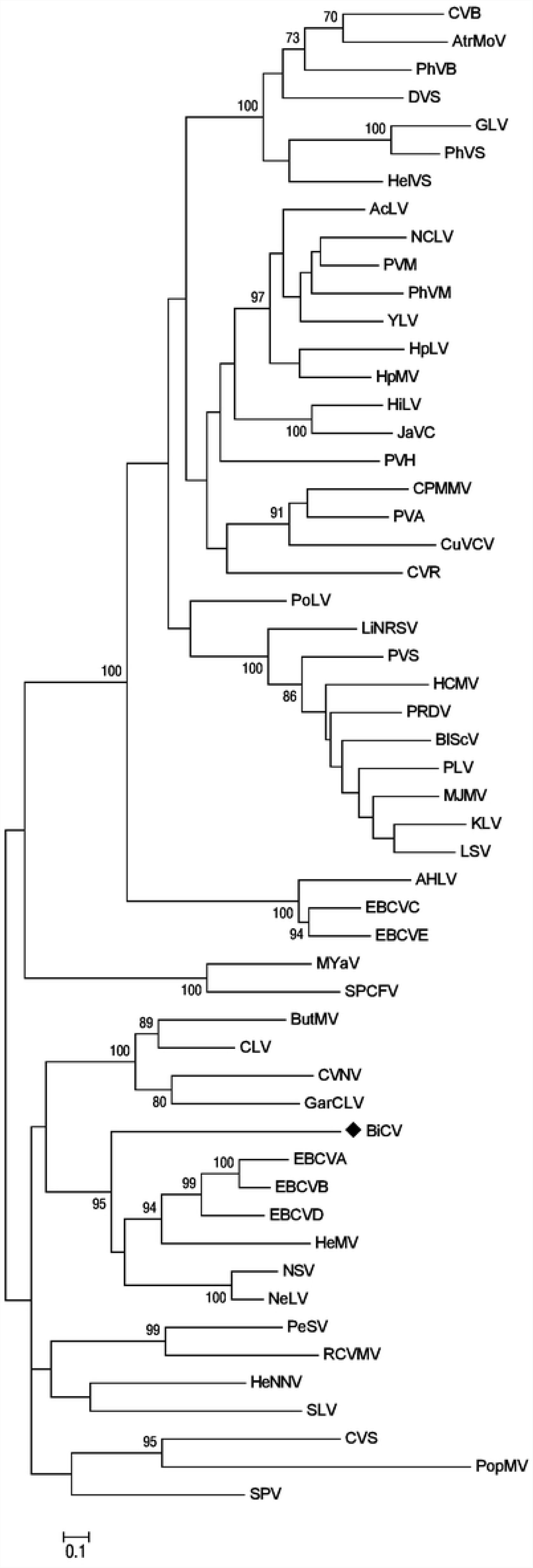
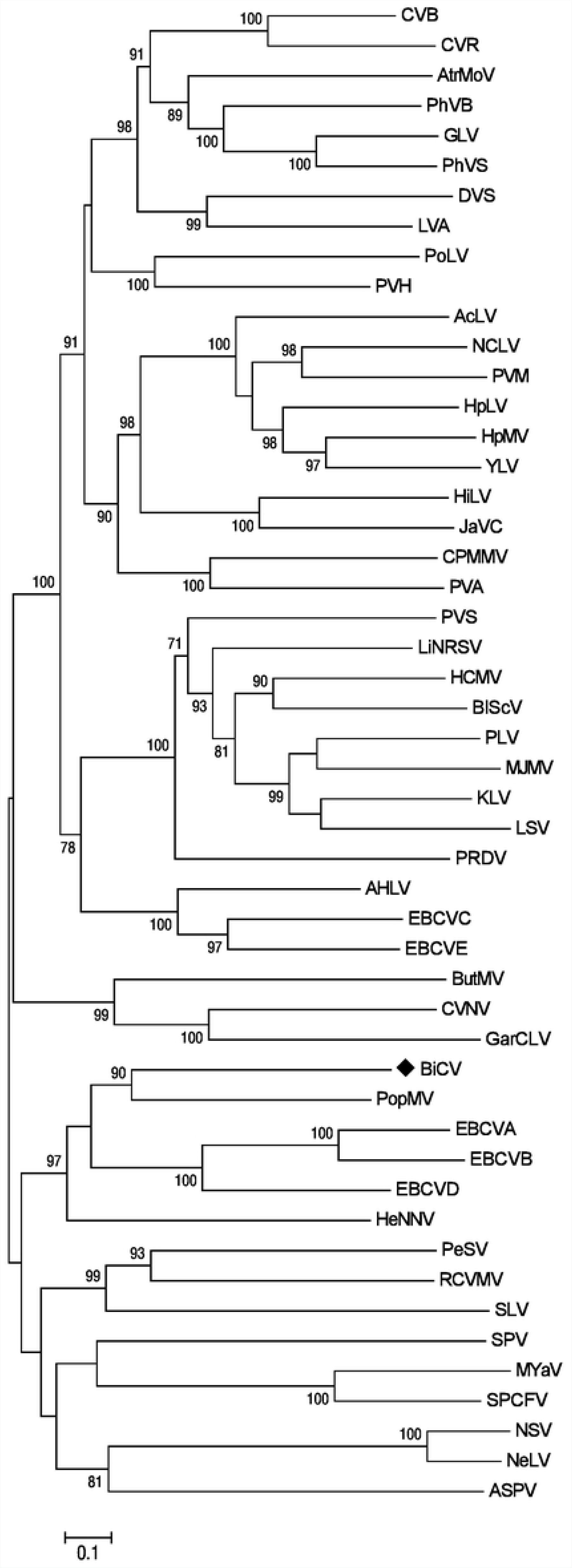
Phylogenetic trees reconstructed using the amino acid sequences of the RdRp (A) and the CP (B) of carlaviruses. The tree was reconstructed using the Maximum Likelihood method and the statistical significance of branches was evaluated by bootstrap analysis (1,000 replicates). Only bootstrap values above 70% are indicated. The scale bar represents 10% amino acid divergence. Virus abbreviations and accession numbers are as follows: aconitum latent virus (AcLV, NC_002795.1), american hop latent virus (AHLV, NC_017859.1), apple stem pitting virus (ASPV, NC_003462.2), atractylodes mottle virus (AtrMoV, KR349343.1), birch carlavirus (BiCV), blueberry scorch virus (BlScV, NC_003499.1), butterbur mosaic virus (ButMV, NC_013527.1), carnation latent virus (CLV, AJ010697.1), carrot virus S (CVS, EU881919), chrysanthemum virus B (CVB, NC_009087.2), chrysanthemum virus R (CVR, MG432107.1), coleus vein necrosis virus (CVNV, NC_009764.1), cowpea mild mottle virus (CPMMV, NC_014730.1), cucumber vein-clearing virus (CuVCV, JN591720.1), daphne virus S (DVS, NC_008020.1), elderberry carlavirus A (EBCVA, NC_029085.1), elderberry carlavirus B (EBCVB, NC_029086.1), elderberry carlavirus C (EBCVC, NC_029087.1), elderberry carlavirus D (EBCVD, NC_029088.1), elderberry carlavirus E (EBCVE, NC_029089.1), gaillardia latent virus (GLV, NC_023892.1), garlic common latent virus (GarCLV, NC_016440.1), helenium virus S (HelVS, D10454.1), helleborus mosaic virus (HeMV, FJ196838.1), helleborus net necrosis virus (HeNNV, NC_012038.1) hippeastrum latent virus (HiLV, NC_011540.1), hop latent virus (HpLV, NC_002552.1); hop mosaic virus (HpMV, NC_010538.1), hydrangea chlorotic mottle virus (HCMV, NC_012869.1), jasmine virus C (JaVC, NC_030926.1), kalanchoe latent virus (KLV, NC_013006.1), ligustrum necrotic ringspot virus (LiNRSV, NC_010305.1), ligustrum virus A (LVA, NC_031089.1), lily symptomless virus (LSV, NC_005138.1), melon yellowing-associated virus (MYaV, LC224308.1), mirabilis jalapa mottle virus (MJMV, NC_016080.1) narcissus common latent virus (NCLV, NC_008266.1), narcissus symptomless virus (NSV, NC_008552.1), nerine latent virus (NeLV, NC_028111.1), passiflora latent virus (PLV, NC_008292.1), pea streak virus (PeSV, NC_027527.1), pepper virus A (PVA, NC_034376.1), phlox virus B (PhVB, NC_009991.1), phlox virus M (PhVM, FJ159381.1), phlox virus S (PhVS, NC_009383.1), poplar mosaic virus (PopMV, NC_005343.1), potato latent virus (PoLV, NC_011525.1), potato virus H (PVH, NC_018175.1), potato virus M (PVM, NC_001361.2), potato virus S (PVS, NC_007289.1), potato rough dwarf virus (PRDV, NC_009759.1), red clover vein mosaic virus (RCVMV, NC_012210.1), shallot latent virus (SLV, NC_003557.1), sweet potato chlorotic fleck virus (SPCFV, NC_006550.1), sweet potato virus (SPV, NC_018448.1), yam latent virus (YLV, NC_026248.1).

## Discussion

The first results of the birch virome characterization presented here occurred after long-term trials aiming to define the causal agent of the “birch leaf-roll disease” which seriously affects birch forests and urban greens throughout Europe. As expected, the development of an HTS strategy succeeded in offering information about the entire virome of the analysed trees. In the case of the symptomatic *B. pendula* sample, the virome comprises five viral agents, namely a new isolate from the well-characterized CLRV nepovirus, two variants of the recently discovered birch leafroll-associated badnavirus (BLRaV), an isolate of the newly discovered birch carlavirus, as well as a partial sequence of the novel birch capillovirus (BCV) (Fig 2; C). In the case of the symptomatic *B. pubescens* seedlings, the virome is less complex; in BpubFinn407501-3A a BLRaV and a BCV infection were detected (Fig 2; A), while in BpubFinn407507_3I a single BLRaV-infection was detected (Fig 2; B). In earlier studies, both *B. pubescens* seedlings were tested CLRV-positive in RT-PCR assays [2]. However, CLRV was not detectable in the samples at the time of the season when the RNA-Seq was performed. This is not surprising according to our experience, because the low CLRV titers in Finnish birches often lead to false negatives when performing diagnostic RT-PCRs [2].

The virome of these three symptomatic birch seedlings analysed by RNA-Seq allows a new interpretation concerning the BLRD etiology. BLRaV should be considered as the main agent directly related with the symptoms, because it is the only virus systematically detected by HTS in symptomatic trees and because it was absent from symptomless ones. Concretely, in the case of BpubFinn407507_3I, it is the only virus with significant number of reads, which tends to demonstrate its causal role. Concerning BIV, it was detected in a symptomless tree, it is therefore, possible that this virus is latent. The other novel virus, BiCV, is detected in only one from the three symptomatic trees, suggesting that most probably it is not needed for BLRD development. BCV was detected in symptomatic and non-symptomatic trees and is thus most possibly a latent virus, although it could still possibly contribute to pathogeny through interactions with other agents.

The complexity of viromes observed in the tested birch seedlings has been lately observed in other birches in Berlin. In an investigation of viruses’ distribution in urban parks in Berlin for two consecutive years, BiCV was present in 16% of the tested birches [26]. In five from these trees a co-infection of BiCV, CLRV and the BLRaV badnavirus was found (data not shown). Co-infection with BiCV, CLRV, ApMV (*Apple mosaic virus*) and BLRaV in birches was observed in urban green of Berlin with an incidence in symptomatic leaves of around 29 % within three years of investigation. These data indicate that mixed infections in birch are widespread A correlation of this viral complex with specific symptom appearance and differentiation of symptomatology in cases of infection by a single virus or by two virus species is not easy to establish [27]. In contrast to annual plants, in a large-volume birch tree the symptomatology may differ in different parts of the canopy and this may be related to differentiation of the virus population – as recently demonstrated in birches [2] - or to other parameters.

Under the light of a holistic understanding of the disease pathogenesis the “pathobiome” concept has been developed, which represents the pathogenic agent integrated within its biotic environment [28]. Understanding the pathobiome thus requires (1) an accurate knowledge of the microorganism community, (2) clear evidence of any effect(s) this microorganism community has on pathogenesis, (3) an understanding of the impact of the microorganism community on persistence, transmission and evolution of pathogenic agents, and (4) knowledge of biotic and abiotic factors that may disrupt the pathobiome and lead to onset of pathogenesis. According to this concept the diverse viruses detected in birches in the present study may play a direct or indirect role in disease development, as each virus may interact with or disturb the virome, ultimately causing a disease [22].

Apart from the virome extensively described here, attention should be also given to the rest of microorganisms detected in the samples. Bacterial species (*Proteobacteria*, *Terrabacteria*, FCB Group bacteria), unclassified Totiviruses as well as thousands of unassigned reads are part of the birch metagenome. Our findings should be examined under the holistic view of the “hologenome theory” [29], which proposes that plants must not be viewed as autonomous entities but rather as holobionts, within which all interacting organisms contribute to the overall stability of the system [30]. Driving factors as microbiota in the soil, the rhizosphere, the rhizoplane, the endosphere and the aboveground compartment play significant role in the health status of the holobiont [31]. With our study we provide some new data regarding the birch microbiote complexity. Their role is not analysed in the present study, but it may be combined with further data in the future.

Concerning the new capillovirus BCV, given the short length of the genomic region characterized, there are still some doubts whether the detected sequences indeed represent an existing virus and, if so, whether this virus can unambiguously be assigned to the Capillovirus genus. This can only be sorted out, ultimately, by efforts to obtain the full genome of the suspected virus. It is noteworthy, that a 600-nt sequence with very high homology with the newly discovered contig has been identified within the transcriptomic data generated from pollen of *Betula verrucosa* [32], indicating that if indeed the sequence identified here is viral, the agent might be more broadly present in other *Betula* species.

To our knowledge, it is the first time that metagenome data of a forest tree species (*Betula sp*.) are reported. In comparison to cultivated plants, little steps are done regarding knowledge on viruses present in forest ecosystems. Missing data or unawareness concerning viral incidence in forests may lead to unjustified disease diagnosis and determination of the causal agent. Not only birch suffers from viral diseases. Virus-like symptoms are commonly observed in *Fraxinus* sp. (Central Europe, Switzerland, Germany), in *Quercus* sp. (Germany, Sweden, Romania), in *Ulmus* sp. (Germany, Sweden, Gotland), in *Acer* sp. (Germany), in *Populus* sp. (Germany, Finland), in Ulmus sp. (Germany), and in *Sorbus* sp. (Germany, North and Central Europe) [2]. Based on our extended experience on recognising symptomatology of viral causal agents and on monitoring distribution of viral diseases, we suggest that viral infections alter plant predisposition and do have an impact on the disease status of many forest and urban trees. HTS technologies may offer a deeper investigation of the viruses in forest species and fill in the knowledge gap concerning the virome of a forest. Current [33] and future investigations are expected to enlighten interacting potential of viruses with influencing abiotic and biotic factors in forest and urban trees as well as the mode of virus transmission.

## Accession numbers

MK402281: Cherry leaf roll virus, complete genome, RNA1

MK402282: Cherry leaf roll virus, complete genome, RNA2

MK402235: Birch idaeovirus, partial genome, RNA1

MK402236: Birch idaeovirus, partial genome, RNA2

MK402233: Birch capillovirus, partial sequence, isolate BpenGer407526_B5

MK402234: Birch capillovirus, partial sequence, isolate BpubFinn407501_3A

## Acknowledgments

This work is partly accomplished in the frame of the COST Action FA1407 with the title ‘Application of next generation sequencing for the study and diagnosis of plant viral diseases in agriculture’. Moreover, we appreciate DFG (Deutsch Forschunsgessellschaft) for funding the projects BU890/14-1 (“Modes of transmission of *Cherry leaf roll virus*”) and BU890/23-1 (“Modes of vector transmission of CLRV – molecular basis and potential arthropod vector species”), through which the scientific basis for accomplishing the given study was built.

## Supporting information

**Fig 7. Phylogenetic trees reconstructed using the amino acid sequences of the proteins encoded by the TGBP1 (A), the TGBP2 (B), the TGBP3 and the ORF6 of carlaviruses.** The trees were reconstructed using the Maximum Likelihood method and the statistical significance of branches was evaluated by bootstrap analysis (1,000 replicates). Only bootstrap values above 70% are indicated. The scale bar represents 10% amino acid divergence.

